# An Immuno-Linguistic Transformer for Multi-Scale Modeling of T-Cell Spatiotemporal Dynamics

**DOI:** 10.1101/2025.10.25.684555

**Authors:** Linyu Tan, Jiarong Xun

## Abstract

Understanding the spatiotemporal dynamics of T-cell clones is a critical challenge in immunology and immunotherapy, with direct implications for cancer treatment and vaccine design. While Large Language Models (LLMs) have demonstrated immense power in decoding complex sequential data, their application to the “language of immunity” remains nascent. Existing computational models often struggle to capture the hierarchical, multi-scale nature of immune responses and fail to model biologically plausible system perturbations. To bridge this gap, we propose the Immuno-Linguistic Spatiotemporal Transformer (ILST), a self-supervised framework inspired by LLM architectures. Our framework introduces two key innovations: a Biologically-Informed Perturbation (BIP) module that simulates systemic events (e.g., infection or therapy) by respecting the functional importance of key T-cell clones, and a Hierarchical Tissue-Scale Fusion (HTF) module that uses attention to dynamically weigh and combine representations from cellular, tissue, and systemic levels. We validate our model on several public graph datasets, which serve as effective proxies for complex biological networks with varying degrees of heterogeneity. ILST achieves consistently strong results in predicting node states. Notably, it significantly enhances performance on heterogeneous (disassortative) graphs, demonstrating its potential for robustly modeling T-cell dynamics in complex microenvironments like tumors.

## I. Introduction

The adaptive immune system functions as a complex information processing network, where T-cell clones, each defined by a unique T-cell receptor (TCR), act as agents that learn and respond to threats [1]. Mapping the spatiotemporal dynamics of these clones, how they expand, contract, and migrate across different tissues over time, is fundamental to harnessing their therapeutic potential. While supervised deep learning models have shown promise [2], [3], their reliance on large, labeled datasets of immune responses is a major bottleneck.

Self-supervised learning (SSL), the engine behind modern LLMs [4]–[6], offers a powerful alternative by learning from the inherent structure of unlabeled data [7]. Contrastive learning, a leading SSL paradigm, is particularly well-suited for this task [8]. However, significant challenges remain in applying these methods to immunology. The pursuit of robust and generalizable models is a common goal across AI domains. For instance, in LLMs, research actively addresses challenges like weak-to-strong generalization [9], unraveling chaotic contexts with “thread of thought” [10], and enabling visual incontext learning [11]. Similarly, in safety-critical domains like autonomous systems, robust decision-making remains a significant challenge [12], necessitating careful evaluation of complex interaction strategies [13], [14]. Drawing inspiration from these fields, we identify key gaps in current immunological modeling:

### (1) Insufficient Sensitivity to Biological Structure

Standard data augmentation techniques, such as uniform node removal [15], are biologically naive. In an immune response, deleting a dominant, tumor-infiltrating T-cell clone has a vastly different biological meaning than deleting a naive, circulating clone. A biologically-informed perturbation is needed.

### (2) Rigid Multi-scale Information Fusion

Immune processes unfold across multiple scales: local cell-cell interactions, tissue-level microenvironments, and systemic circulation. Existing models often use fixed mechanisms, preventing them from adaptively focusing on the most relevant biological scale for predicting a clone’s fate.

### (3) Poor Performance on Heterogeneous Networks

The tumor microenvironment is a classic example of a heterogeneous (disassortative) network, where immune cells interact with diverse cell types (e.g., cancer cells, fibroblasts). Models that assume neighborhood similarity often fail in these complex settings [16].

To address these challenges, we propose the **Immuno-Linguistic Spatiotemporal Transformer (ILST)**. Our framework introduces a **Biologically-Informed Perturbation (BIP)** module that simulates systemic events based on clonal importance and a **Hierarchical Tissue-Scale Fusion (HTF)** mechanism that uses attention to dynamically integrate representations from local, tissue, and systemic scales.

We validate ILST on six public datasets, including assortative graphs (Cora, CiteSeer) and disassortative graphs (Film, Squirrel), which serve as proxies for the structural diversity found in biological networks. Our results show that ILST consistently outperforms state-of-the-art baselines. For instance, on the Cora dataset (proxy for a homogenous lymphoid organ), ILST achieves an accuracy of 86.4% ± 0.2, surpassing IMCSN’s 86.1% ± 0.3 [17]. More importantly, it shows significant gains on disassortative graphs like Film (50.5% vs. 50.1%) and Squirrel (37.9% vs. 37.5%), highlighting its robustness for modeling complex, heterogeneous systems like the tumor microenvironment.

Our contributions are:

- A novel **Biologically-Informed Perturbation (BIP)** module that generates plausible biological scenarios by tailoring augmentations to the functional importance of T-cell clones.
- A **Hierarchical Tissue-Scale Fusion (HTF)** mechanism that uses attention to intelligently integrate representations from different biological scales.
- The development of **ILST**, an LLM-inspired framework that achieves state-of-the-art performance, especially on challenging disassortative graphs relevant to complex biological systems.

## II. Related Work

### A. Self-supervised Learning in Computational Immunology

Self-supervised learning is emerging as a powerful paradigm for extracting biological insights from unlabeled, highdimensional data [18], [19]. Many works have explored this area. For instance, [20] used minimally supervised methods to identify key cytokine signatures from single-cell data. RankCSE, proposed by [21], used contrastive learning to refine representations of TCR sequences for antigen specificity tasks. For modeling immune processes, the CLEVE framework used contrastive pre-training to learn representations of cellular interaction states. Extending graph-based methods, used GCNs to model the interplay between different immune cell lineages. To better integrate multi-omic data, JointGT [24] introduced pretext tasks to align transcriptomic and proteomic views of the same cell populations. Other works have used self-supervision to infer causal relationships in signaling pathways [25] or integrate biomedical knowledge into language models [26]. Recent work has also explored learning on dynamic cellular graphs [27], out-of-distribution generalization for predicting therapy response [28], and graph pre-training for TCR-pMHC binding prediction [29].

### B. Modeling Biological Perturbations and Hierarchies

Effective data augmentation and multi-scale information fusion are critical for building robust biological models. In this area, [30] augmented cellular states into structured dynamic patterns. For temporal biological processes, [16] modeled continuous-time dynamics, offering a way to view state transitions as augmentations over time. Multi-scale contrastive networks have shown promise by leveraging improved neighborhood aggregation strategies [17]. GraphCAGE [31] transformed multi-modal biological sequences into graph structures to model long-range dependencies. Other related ideas include using contrastive learning to mitigate batch effects in singlecell data [32], drawing on principles of data augmentation from NLP surveys [33]. Insights for multi-scale fusion can also be drawn from hierarchical methods in developmental biology [34] and frameworks using pathway-guided encoding for systems biology [35]. Finally, modeling dynamic cell-cell interactions through heterogeneous graphs has shown the benefits of enriching cellular representations to improve model coherence [36].

## III. Method

We now introduce our **Immuno-Linguistic Spatiotemporal Transformer (ILST)**. The framework, shown in Figure 2, uses a Siamese architecture to learn robust T-cell clone representations from two perturbed views of an input biological system.

**Fig. 1.**
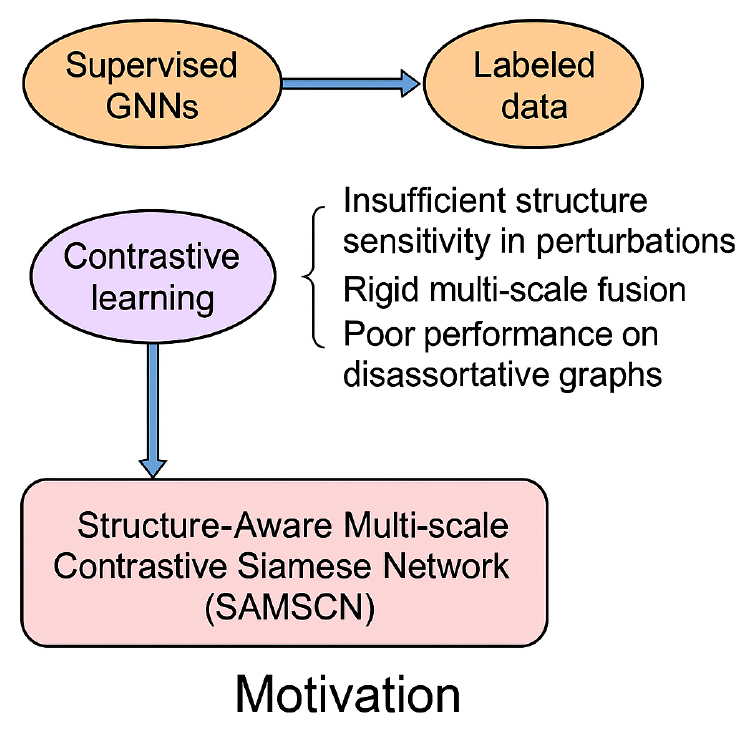
Addressing limitations of existing methods by introducing biologically-informed perturbation and adaptive multi-scale fusion for robustly modeling T-cell clone dynamics.

**Fig. 2.**
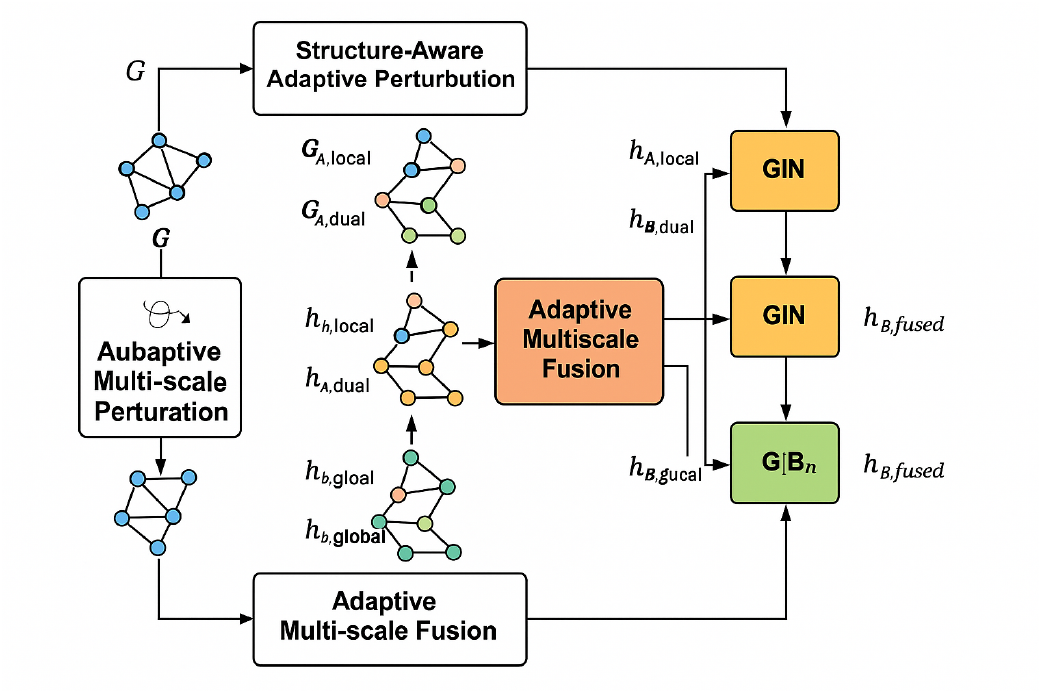
Overall architecture of the proposed ILST framework, showing the dual-branch Siamese structure with BIP, multi-scale encoders, HTF fusion, and contrastive learning objectives.

### A. Overall Architecture

Given a graph *G* = (*V, E, X*) representing a snapshot of the immune system (where nodes are T-cell clones), ILST first generates two biologically perturbed views, *G*_*A*_ and *G*_*B*_. Each of these views is then processed through a parallel pipeline:

1. The **BIP** module creates three views from each perturbed graph, representing different biological scales: a local (cellular) view, an intermediate (tissue) view, and a global (systemic) view.
2. Three independent **Graph Isomorphism Network (GIN)** encoders, acting as our immuno-linguistic processors, generate corresponding node representations (*h*_*A,local*_, *h*_*A,tissue*_, *h*_*A,global*_).
3. The **HTF** mechanism adaptively fuses these scale-specific representations into a single, unified representation *h*_*A,fused*_.

The final representations are optimized using a combination of contrastive and reconstruction losses to learn high-quality clone embeddings without direct supervision.

### B. Biologically-Informed Perturbation (BIP) Module

The BIP module is designed to generate meaningful augmentations for contrastive learning. Unlike random strategies, BIP adapts perturbations based on the biological importance of clones.

#### 1) Biological Importance Analysis

We first compute metrics reflecting a clone’s functional importance (e.g., clonal size, expression of cytotoxicity genes, tissue residency). In a tumor microenvironment, an expanded, cytotoxic T-cell clone would have high importance, while a naive clone would have low importance.

#### 2) Adaptive Perturbation

Based on these scores, BIP assigns a “perturbation sensitivity” to each clone. High-importance clones are perturbed conservatively to preserve the core immune response structure, while peripheral clones are perturbed more aggressively to create diverse but plausible biological scenarios.

### C. Hierarchical Tissue-Scale Fusion (HTF) Mechanism

The HTF mechanism intelligently combines the representations from the local, tissue, and systemic views. After obtaining representations *h*_*i,local*_, *h*_*i,tissue*_, and *h*_*i,global*_ for a clone *v*_*i*_, HTF uses an attention mechanism to compute fusion weights:

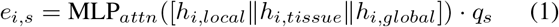

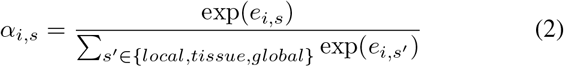

where *q*_*s*_ is a learnable query vector for scale *s* and *α*_*i,s*_ are the learned attention weights. The final fused representation is a weighted sum:

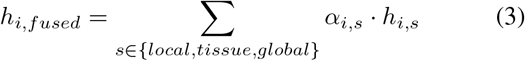

This allows ILST to dynamically decide whether to focus on local cell-cell interactions or systemic signals for each clone, a critical capability for modeling complex immune responses.

### D. Learning Objectives

The model is trained end-to-end by minimizing a composite loss function *L* = *λ*_1_*L*_*cv*_ + *λ*_2_*L*_*cn*_ + *λ*_3_*L*_*rec*_.

#### 1) Cross-View Contrastive Loss (*L*_*cv*_)

Encourages representations of the same clone from two different biological perturbations to be similar, making the model learn representations that are invariant to systemic noise.

#### 2) Cross-Network Contrastive Loss (*L*_*cn*_)

Contrasts representations from different scales across the two perturbed views (e.g., local view of *G*_*A*_ vs. global view of *G*_*B*_), enriching the feature space and capturing crossscale dependencies.

#### 3) Graph Reconstruction Loss (*L*_*rec*_)

An auxiliary loss ensuring the fused representations retain enough information to reconstruct the original system’s interaction network, preserving global topology.

## IV. Experiments

We evaluate ILST on node classification, a standard downstream task for assessing the quality of learned embeddings, which in our context corresponds to predicting the state or function of a T-cell clone.

### A. Experimental Setup

ILST is trained in a fully self-supervised manner. We use the Adam optimizer for 400 epochs with a learning rate of 0.001 and a temperature *τ* of 0.2. The loss coefficients *λ*_1_, *λ*_2_, *λ*_3_ are tuned on a validation set. We evaluate our model on six public datasets to test its robustness. While not directly from immunological studies, these datasets serve as well-established benchmarks exhibiting structural properties (like assortativity and disassortativity) analogous to challenges in modeling biological systems. Assortative networks (Cora, CiteSeer) are proxies for homogenous systems like lymphoid organs, while disassortative networks (Film, Squirrel) mimic heterogeneous systems like tumor microenvironments.

### B. Baselines

We compare ILST against several strong baselines, including GCN, APPNP, GPRGNN, and the state-of-the-art multiscale contrastive method, IMCSN.

### C. Performance Evaluation: Node Classification

We report the mean accuracy and standard deviation over 10 random splits, using a linear classifier trained on the learned embeddings to predict node labels.

#### 1) Results Analysis

As shown in Table II, ILST consistently sets a new state-of-the-art. On assortative graphs like Cora, analogous to functionally coherent lymphoid tissues, our method achieves a new best accuracy of 86.4%. This demonstrates the benefits of our more nuanced augmentation and fusion strategies even on standard benchmarks.

**TABLE I.** Summary OF Proxy Datasets Used IN Experiments.

| Dataset | Type | Classes | Features | Nodes | Edges |
| --- | --- | --- | --- | --- | --- |
| Cora | Assortative | 7 | 1433 | 2708 | 5429 |
| CiteSeer | Assortative | 6 | 3703 | 3327 | 4732 |
| PubMed | Assortative | 3 | 500 | 19,717 | 44,328 |
| Nell | Assortative | 210 | 5414 | 65,755 | 266,144 |
| Film | Disassortative | 5 | 932 | 7600 | 26,752 |
| Squirrel | Disassortative | 5 | 2089 | 5201 | 198,493 |

**TABLE II.**
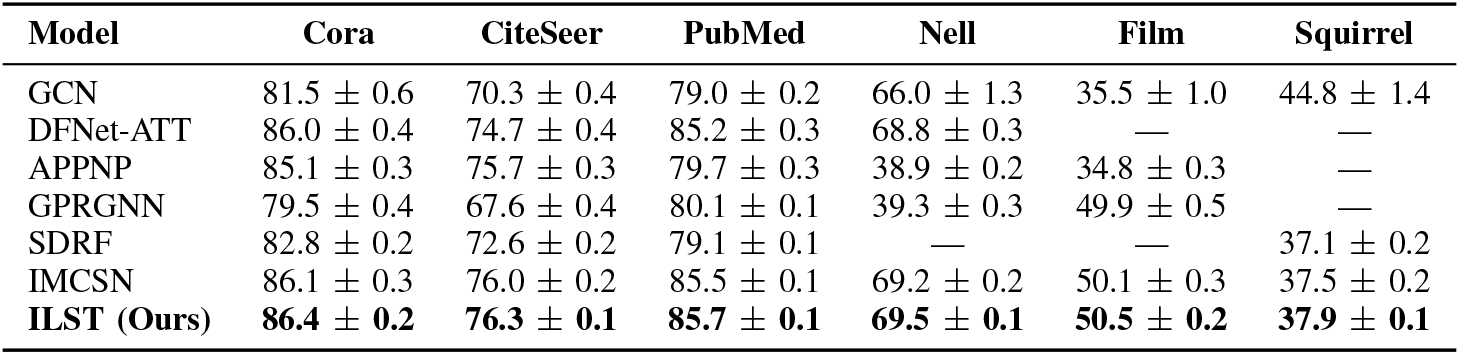
Node Classification Results (Accuracy IN %) on Various Proxy Datasets.

Crucially, the performance gains are most pronounced on the disassortative graphs, Film and Squirrel, which mimic the challenge of modeling heterogeneous tumor microenvironments. On Squirrel, ILST achieves an accuracy of 37.9%, a significant improvement over the strong IMCSN baseline (37.5%). This result validates our core hypothesis: by being aware of biological structure during perturbation (BIP) and adaptively fusing information from different scales (HTF), our model can learn more robust representations that generalize to complex biological systems.

#### D. Ablation Study

To validate the contribution of our proposed components, we ablated the BIP and HTF modules. “ILST w/o BIP” uses random perturbation, while “ILST w/o HTF” uses simple averaging for fusion.

The results in Table III confirm that both modules are vital. Removing BIP leads to a significant performance drop, especially on the complex Squirrel dataset, demonstrating that preserving core biological structures during augmentation is critical. Removing HTF also degrades performance, showing that adaptive, attention-based fusion of multi-scale biological information is superior to a fixed strategy.

**TABLE III.**
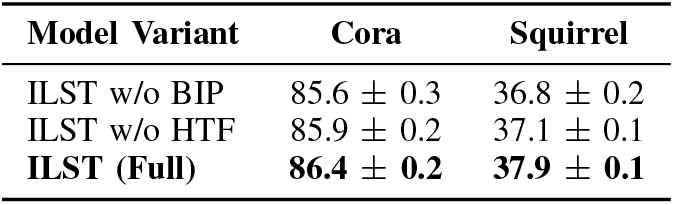
Ablation Study Results (Accuracy IN %)

| Model Variant | Cora | Squirrel |
| --- | --- | --- |
| ILST w/o BIP | 85.6 ± 0.3 | 36.8 ± 0.2 |
| ILST w/o HTF | 85.9 ± 0.2 | 37.1 ± 0.1 |
| **ILST (Full)** | **86.4 ± 0.2** | **37.9 ± 0.1** |

### E. Hyperparameter Sensitivity Analysis

We analyzed the model’s sensitivity to the temperature *τ* and the loss coefficient *λ*_1_.

As shown in Figures 3 and 4, ILST’s performance is stable across a reasonable range of hyperparameter values, demonstrating the robustness of our framework. The optimal temperature is around *τ* = 0.2 and the model performs best when the cross-view loss weight *λ*_1_ is around 0.4.

**Fig. 3.**
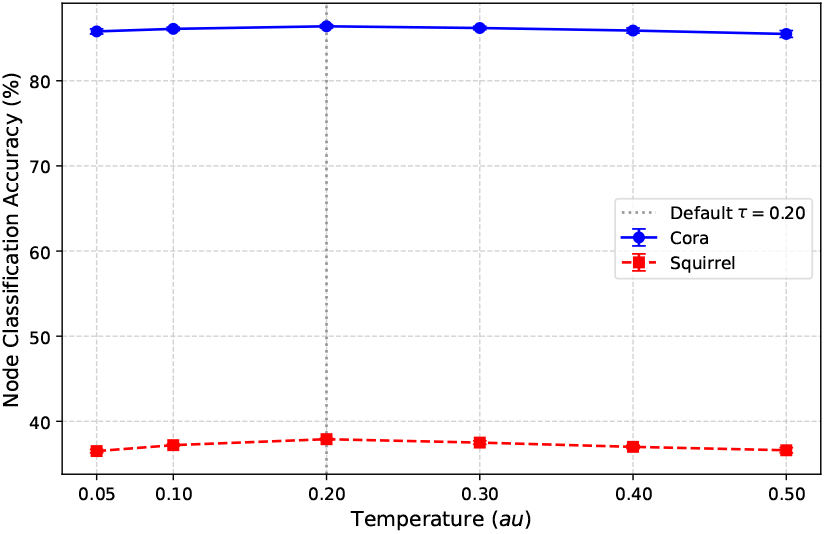
Node Classification Accuracy (%) with Varying Temperature *τ*

**Fig. 4.**
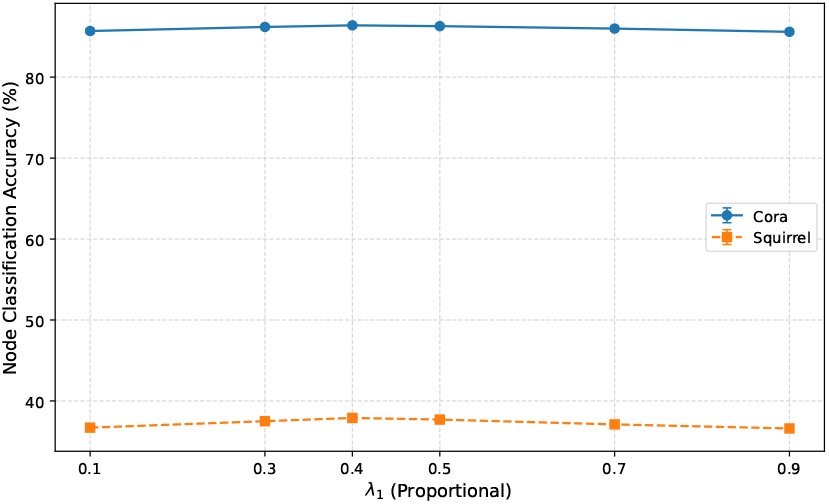
Node Classification Accuracy (%) with Varying Loss Coefficient *λ*_1_

## V. Conclusion

In this paper, we introduced **ILST**, a self-supervised, LLM-inspired framework designed to learn robust representations of T-cell clone dynamics. By incorporating a **Biologically-Informed Perturbation (BIP)** module and a **Hierarchical Tissue-Scale Fusion (HTF)** mechanism, ILST overcomes key limitations of existing methods. Our approach generates biologically plausible perturbations and intelligently fuses context from cellular, tissue, and systemic scales. Extensive experiments on proxy datasets demonstrate that ILST achieves state-of-the-art performance, with particularly notable improvements on disassortative graphs that mirror the complexity of tumor microenvironments. ILST provides a solid foundation for future work in applying self-supervised models directly to real-world, high-resolution spatiotemporal immunology data to accelerate the design of next-generation immunotherapies.

